# Prediction of Resistant Mutations against Upcoming ALK-TKIs, Repotrectinib (TPX-0005) and Ensartinib (X-396)

**DOI:** 10.1101/2022.05.26.493531

**Authors:** Yuta Doi, Hiroaki Tagaya, Ayaka Noge, Kentaro Semba

## Abstract

ALK gene rearrangement is observed in approximately 4% of patients with non-small cell lung cancer. These individuals benefit clinically from a range of approved ALK-TKIs; however, using many ALK-TKIs in a row will always result in resistant compound mutations in the kinase domain. Therefore, next-generation ALK-TKIs which are potent to these resistant mutations are still under development. In this context, preclinical prediction of resistant mutations generated by developing ALK-TKIs will provide useful information about the effective sequential treatment of ALK-TKIs. In this study, we developed a simple error-prone PCR-based mutation prediction system, and as a model study, we predicted resistant mutations against upcoming ALK-TKI, repotrectinib, and ensartinib. According to the predictive mutation patterns, repotrectinib and ensartinib may be used as second-line therapeutic options following the first-line alectinib treatment.

## Introduction

In about 4% of patients with non-small cell lung cancer (NSCLC), chromosomal rearrangements involving the anaplastic lymphoma kinase (ALK) gene are found^1,2^. ALK rearrangement results in constitutive activation of ALK due to self-oligomerization through fusion partners such as echinoderm microtubule-associated protein-like 4 (EML4)^3–5^, and aberrant cancer cell proliferation via ALK downstream pathways^6^.

For the treatment of ALK-positive NSCLC, various ALK tyrosine kinase inhibitors (ALK-TKIs) have been developed. Crizotinib is the first ALK-TKI to be approved for the treatment of NSCLC patients who present with ALK mutations. A phase III trial showed significantly longer progression-free survival and a higher response rate with crizotinib than with standard chemotherapy^7^. In phase III clinical trial, the second-generation ALK-TKI, alectinib, outperformed crizotinib in terms of therapeutic advantages; hence, it is recommended as the first-line treatment for NSCLC. However, patients treated with alectinib relapse within several years mainly due to a secondary point mutation such as G1202R or I1171N in ALK kinase domain^9^. A third-generation ALK-TKI, lorlatinib, was designed to combat these resistant mutations^10,11^, which is currently recommended for the second-line treatment. Because of additional point mutations such as L1196M + G1202R, tumor recurrence is common even after lorlatinib treatment^12,13^. Although some of such compound mutants demonstrate re-sensitivity to other TKIs, no effective therapeutic strategies have been approved against lorlatinib-resistant compound mutants^14–16^. Furthermore, in the CROWN study^17^, lorlatinib showed distinct side effects such as hypercholesterolemia, hypertriglyceridemia, peripheral neuropathy, and cognitive impairments, which may reduce patients’ quality of life. Therefore, lorlatinib may exhibit fewer therapeutic benefits for some patients. At that point, various therapeutic strategies following the alectinib treatment need to be prepared for optimal usage.

Preclinical prediction of resistant mutations against TKIs and discovery of the potent TKIs to their resistant mutants is one of the practical approaches to develop effective therapeutic strategies. Some groups developed a method for predicting resistant mutations using *N*-ethyl-*N*-nitrosourea (ENU) mutagenesis^13,15,18^; however, ENU mutagenesis exhibited a bias for variations of base substitutions and thereby demonstrated probability that all of resistant mutations observed in the clinic might not be detected. Another disadvantage of ENU mutagenesis is that mutations can be induced in the entire genomic DNA, necessitating the analysis of many resistant clones to assess mutations in the ALK kinase domain.

In this report, we developed a controllable and comprehensive approach for predicting resistant mutations using error-prone PCR. As a model, we predicted resistant compound mutations in hypothetical cases that upcoming ALK-TKIs, repotrectinib (TPX-0005), and ensartinib (X-396) are applied after recurrence with alectinib-resistant mutation, G1202R or I1171N. Analyzing the sensitivity of the predicted mutants to other TKIs, we explain the availability of repotrectinib and ensartinib for therapeutic strategies following the first-line alectinib treatment.

## Materials & Methods

### Cell lines and culture condition

Ba/F3 cells, murine bone marrow-derived pro-B cells were cultured in RPMI-1640 (FUJIFILM Wako, Osaka, Japan) supplemented with 10% fatal bovine serum (FBS) (Nichirei Biosci., Tokyo, Japan), 100 U/mL penicillin (Meiji Seika Pharma, Tokyo, Japan), 100 μg/mL streptomycin (Meiji Seika Pharma), and 10 ng/mL murine interleukin-3 (IL-3) (Pepro Tech, Cranbury, NJ, USA) at 37°C and 5% CO_2_. Platinum-E retroviral packaging cells (Plat-E cells) were cultured in D-MEM (low glucose) (FUJIFILM Wako) supplemented with 10% FBS, 100 U/mL penicillin, and 100 μg/mL streptomycin at 37°C and 5% CO_2_.

### Reagents

Crizotinib (PF-02341066) was supplied by LC laboratories. Alectinib (CH5424802) was supplied by ChemScene LLC. Lorlatinib (PF-06463922), repotrectinib (TPX-0005), and ensartinib (X-396) were purchased from Cayman Chemical. Adaphostin was purchased from Sigma-Aldrich. Gilteritinib (ASP2215) and TPX-0131 was supplied by MedChemExpress. All inhibitors were dissolved in dimethyl sulfoxide (DMSO).

### Establishment of EML4-ALK expressing Ba/F3 cells

For retrovirus production, pMXs-GW-IRES-Puro vector containing cDNA of EML4-ALK variant 1 was transfected into Plat-E cells. After 24 hours of incubation, the medium was changed to RPMI-1640, and the retrovirus was harvested. Ba/F3 cells were then infected with the retrovirus with 8 μg/mL polybrene. After 24 h incubation, retrovirus containing medium was changed to RPMI-1640 supplemented with 0.5 ng/mL IL-3. Following a further 24 h incubation, the culture medium was changed to RPMI-1640 supplemented with 0.05 ng/mL IL-3 and 1 g/mL puromycin (FUJIFILM Wako). Puromycin was applied to Ba/F3 cells for 48 h before establishing EML4-ALK expressing Ba/F3 cells. These EML4-ALK expressing Ba/F3 cells were cultured in RPMI-1640 without IL-3. Base substitutions were induced in the pMXs vector with the PrimeSTAR^®^ Mutagenesis Basal Kit (TaKaRa Bio Inc.) before transfection into Plat-E cells to create ALK-TKIs resistant mutants.

### Cell viability assay

Ba/F3 cells were seeded into 96-well plates (2,000 cells/well), and serially diluted inhibitors were added to the culture. After 72 h, the culture was supplemented with Cell Counting Kit-8 (CCK-8) (DOJINDO). The absorbance at 450 nm was measured after 2 h of incubation, and the data were analyzed using GraphPad Prism version 6 (GraphPad Software).

### Antibodies and immunoblotting

Ba/F3 cells (1 × 10^6^ cells) were seeded into 12-well plates and treated with inhibitors for 3 h. PBS was used to wash the cells, which were then suspended in TNE buffer (10 mM Tris-HCl (pH 7.4), 1 mM EDTA, 150 mM NaCl, and 1% NP-40). Total protein concentration was measured with Pierce™ BCA Protein Assay Kit (ThermoFisher Scientific). Then, the suspension was treated with 2 × sample buffer containing 100 mM Tris-HCl (pH 6.8), 4% sodium dodecyl sulfate (SDS), 20% glycerol, 10% 2-mercaptoethanol, and 0.01% bromophenol blue. The samples were well sonicated and boiled at 95°C for 5 min. Then, the samples’ total proteins were electrophoresed in 7.5% SDS-polyacrylamide gels. The proteins were transferred from the gels to Immobilon PVDF membranes (Merck Millipore Ltd.). The membranes were immersed in appropriate blocking buffer (Tris-buffered saline 0.05% Tween 20 (TBST) supplemented with 5% w/v bovine serum albumin or 5% w/v nonfat dry milk) for 1 h at room temperature. After that, the membranes were incubated overnight at 4°C with gentle agitation in primary antibody dilution buffer: phosphorylated ALK (Y1604) (Cell Signaling Technologies, #3341, 1:1000), ALK (Cell Signaling Technologies, #3791, 1:1000), and α-tubulin (Wako, 013-25033, 1:2000). After washing with TBST, the membranes were incubated for 1 hour at room temperature with gentle agitation in the blocking buffer supplemented with appropriate horseradish peroxidase (HRP)-linked secondary antibodies: anti-rabbit IgG, HRP-linked antibody (Cell Signaling Technologies, #7074, 1:2000) and anti-mouse IgG, HRP-linked antibody (Cell Signaling Technologies, #7076, 1:2000). Then the membranes were washed with TBST and incubated with Immobilon^®^ Western Chemiluminescent HRP Substrate (EMD Millipore Corporation) or ImmunoStar^®^ LD (Wako) at room temperature for 3 min. Proteins were detected using ChemiDoc XRS+ System (Bio-Rad Laboratories).

### Construction of cDNA mutant libraries by error-prone PCR

Mutations in the pMXs vector were induced at random in a 500-bp region encoding the ALK kinase domain (GenBank: AB663645, 1850-2383) with G1202R mutation, I1171N mutation, or no mutations using the GeneMorph II Random Mutagenesis Kit (Agilent Technologies) according to the manufacturer’s instructions. The mutation rate was set to low (0.4 mutations/kb, 17.2 μg of the pMXs vector in 50 μL PCR reaction solution). pMXs vector arms which had the homologous sequences in 40 bp of their terminals to each terminal sequence of error-prone PCR products were created with using KOD FX Neo (TOYOBO). Next, the products were amplified with PrimSTAR^®^ Max DNA Polymerase (TaKaRa bio Inc.). The vector arms and PCR products were combined by incubating them for 1 hour at 50°C in the Gibson Assembly reaction buffer (5% PEG-8000, 100 mM Tris-HCl (pH 7.4), 10 mM MgCl_2_, 10 mM dithiothreitol (DTT), 0.2 mM dNTPs, 1 mM nicotinamide adenine dinucleotide (NAD+), 0.004 U/μL T5 exonuclease (New England BioLabs Inc.), 0.025 U/ μL Phusion DNA polymerase (Finnzymes)). Ethanol precipitation was used to enrich and desalt combined cDNA before it was introduced into electrocompetent DH10B E. coli. After the electroporation, the competent cells were incubated in SOC medium (2% typtone, 0.5% yeast extract, 10 mM NaCl, 2.5 mM KCl, 20 mM MgSO_4_, 20 mM glucose) for 1 h with intense shaking and then seeded on LB agar plates supplemented with 100 mg/L Ampicillin. After 12 h incubation at 37°C, each mutant cDNA library was prepared from approximately 7 × 10^5^ colonies.

### Establish cDNA mutant libraries expressing Ba/F3 cells and drug screening

cDNA mutant libraries were transfected into Plat-E cells. After 24 hours of incubation, the medium was changed to RPMI-1640, and the retrovirus was harvested. Ba/F3 cells (5 × 10^6^ cells/well, 3 × 10^7^ cells in total) were seeded into 12-well plates and mixed with harvested retrovirus supplemented with 8 g/mL polybrene. Ba/F3 cells were centrifuged in the 12-well plates for 1h at 32°C, 900 × g. Then, the culture was transferred to cell culture flasks and incubated at 37°C and 5% CO_2_. As previously stated, cDNA mutant libraries expressing Ba/F3 cells were established. To assess virus infection efficiency, alive cells were monitored with a hemocytometer after 40 h of puromycin addition by staining dead cells with 0.5% trypan blue when noninfected cells were completely dead. The infection efficiency was calculated by drawing their growth curve and predicting the percentage of infected cells against total alive cells prior to puromycin addition.

Ba/F3 cells (10,000 cells) expressing EML4-ALK with G1202R and a random mutation were seeded into a well of 96-well plates and cultured with repotrectinib (1.5 μM) for 2 weeks. Ba/F3 cells (1,000 cells) expressing EML4-ALK with I1171N and a random mutation were seeded into 96-well plates and cultured for 2 weeks with ensartinib (800 nM). Then, the regions encoding the ALK kinase domain were amplified from the resistant mutants using KOD FX Neo (TOYOBO) polymerase, and resistant mutations were predicted using Sanger sequencing.

### Analysis of cDNA libraries with next-generation sequencer (NGS)

To evaluate the variation and frequency of induced mutations, the cDNA mutant libraries were sequenced with NGS. The 1st PCR was carried out with PrimeSTAR^®^ Max DNA polymerase (TaKaRa Bio Inc.), targeting the region where the mutations were induced by the error-prone PCR. Wizard^®^ SV Gel and the PCR Clean-up System were used to clean the PCR products (Promega). Next, 2nd PCR was performed with TaKaRa Ex Taq^®^ Hot Start Version (TaKaRa Bio Inc.). The PCR products were purified with AMPure XP (BECKMAN COULTER). The amplicons were sequenced with MiSeq (2 × 300 bp). To remove low-quality reads, Fastq files were edited with Trimmomatic by performing a sliding window trimming (Qscore < 30, window size = 5)^20^. Then, the paired reads were aligned against the ALK cDNA sequence, which included the G1202R or I1171N mutations, using HISAT-2^21^. Pysamstats were used to count the number of read bases at each position. The pMXs plasmid incuding cDNA of EML4-ALK wild type was sequenced and analyzed in the same way as a negative control.

The mutant variations were observed by adjusting the count data for the negative control count data. First, the count data at the position where mutation frequency was aberrantly high (> 0.05) in the wild-type sample were removed because these mutations were affected by sequencing and PCR errors. Then, the count data from 3545 to 3744 bp were removed because total reads in this region were much lower than in other regions, and the data would be contaminated by errors. Following that, the count data of each mutation pattern at the position was added together and subtracted from the count data of the wild-type sample.

### Prediction of resistant mutations using NGS

Ba/F3 cells expressing cDNA mutant libraries with G1202R, I1171N mutation, or no mutations (3 × 10^6^ in total) were seeded into 25 cm^2^ culture flask and cultured for 1 week with repotrectinib (1.5 μM), ensartinib (800 nM), or alectinib (50 nM). Then, genomic DNA was extracted using the Gene EluteTM Mammalian Genomic DNA Miniprep Kit (Sigma-Aldrich), followed by NGS analysis as previously stated.

## Results

### Sensitivity of Alectinib-resistant mutants to repotrectinib and ensartinib

Alectinib is currently suggested as the first-line therapy in Japan. To apply other TKIs for second-line therapeutic options against recurrence after alectinib treatment, alectinib-resistant mutants must be sensitive to such TKIs. We established Ba/F3 cells expressing EML4-ALK variant 1 with commonly observed alectinib-resistant mutations, G1202R or I1171N, and tested their sensitivity to two upcoming ALK-TKIs, repotrectinib (TPX-0005) and ensartinib (X-396) (Fig.1a and 1b). CCK-8 cell viability assay indicated that the G1202R mutant was sensitive to repotrectinib and resistant to ensartinib, while the I1171N mutant was resistant to repotrectinib and sensitive to ensartinib (Fig.1c–d), consistent with previous reports^22,23^. Immunoblotting revealed that only repotrectinib or ensartinib inhibited phosphorylation of ALK with the G1202R or I1171N mutations respectively (Fig.S1a–b). As a result, we selected these two compounds to be tested as subsequent therapy options after alectinib treatment.

**Fig 1.**
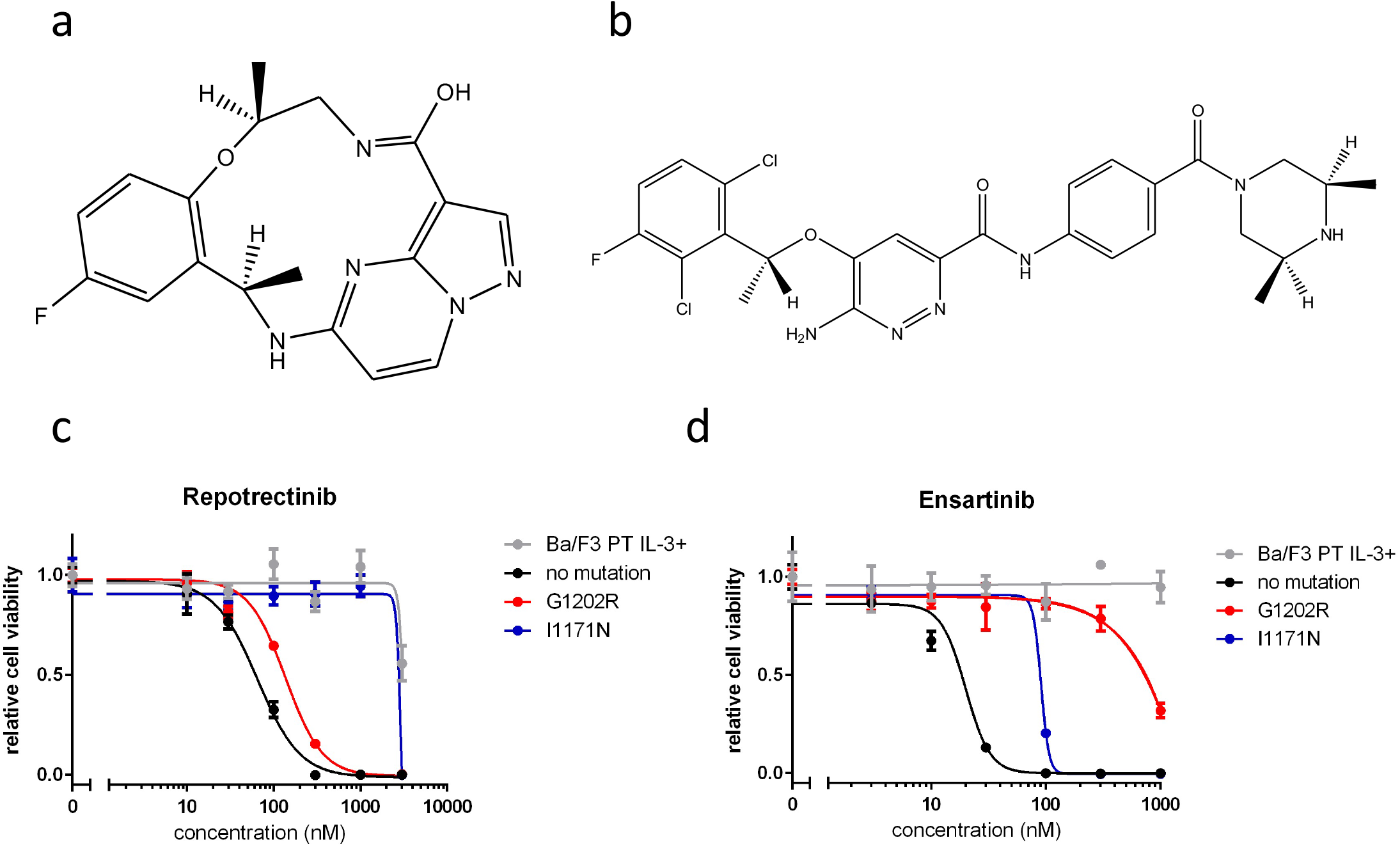
The sensitivity of alectinib-resistant mutants against repotrectinib and ensartinib. (a) Chemical structure of repotrectinib. (b) Chemical structure of ensartinib. (c–d) Sensitivity evaluation of alectinib-resistant mutants against repotrectinib or ensartinib. Ba/F3 cells expressing EML4-ALK variant 1 with G1202R or I1171N were exposed to each inhibitor for 72 h. Cell viability was evaluated by CCK-8 and absorbance at a 450 nm wavelength.

### Experimental system for predicting resistant mutations using error-prone PCR

Since the main cause of recurrence is acquired point mutations in the ALK kinase domain, precise prediction of resistant mutations at the preclinical stage is required to develop effective therapeutic strategies following alectinib treatment. Thus, we established a comprehensive and controllable method using error-prone PCR for preclinical prediction (Fig.2a).

**Fig 2.**
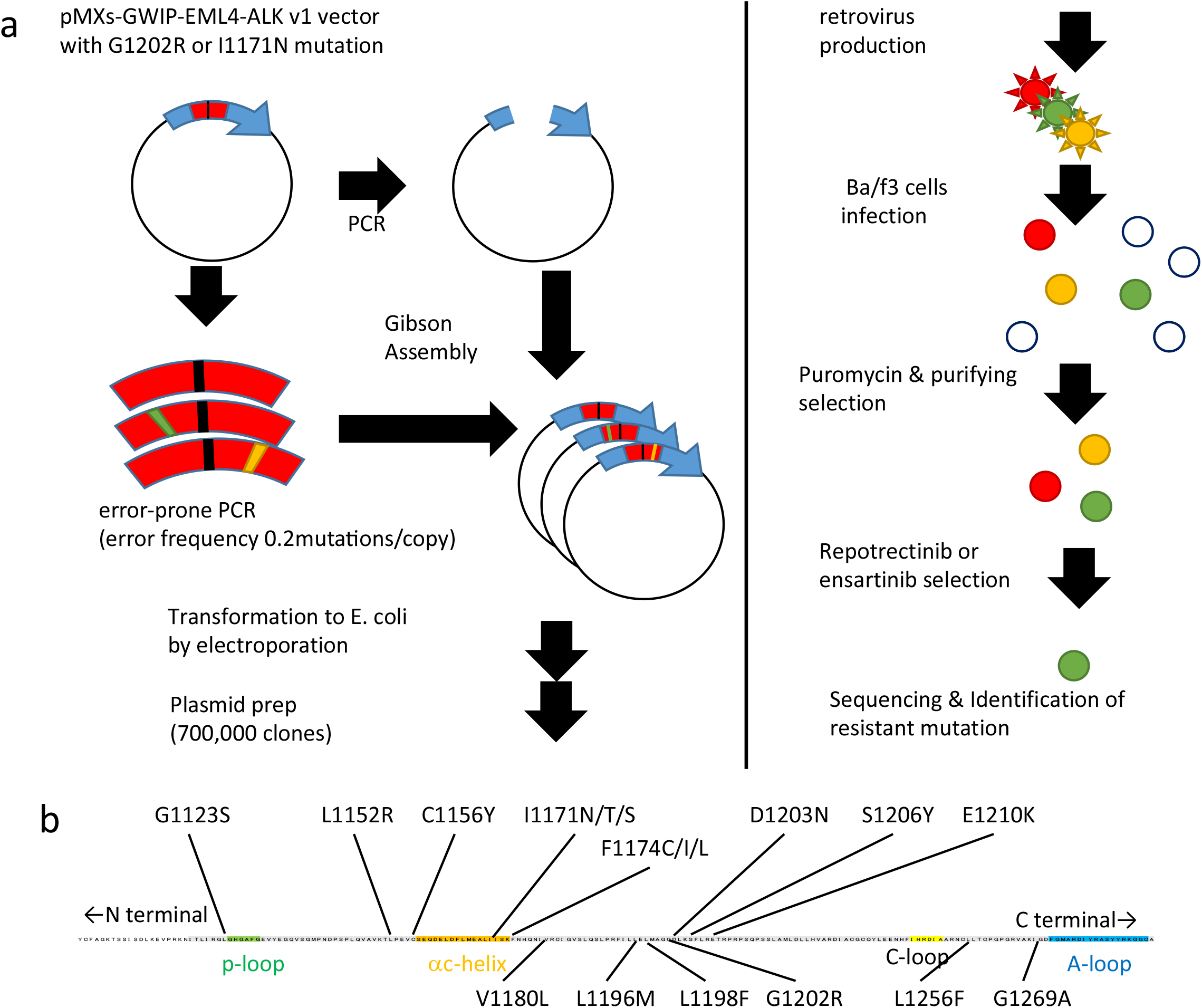
Method for predicting resistant mutations using error-prone PCR. (a) The protocol for predicting resistant mutations using error-prone PCR. Error-prone PCR was used to introduce random mutations into the ALK kinase domain of pMXs-GWIP-EML4-ALK v1 with G1202R or I1171N. PCR products were mixed with vector arms before being transformed into E. coli. cDNA mutant libraries were established by plasmid preparation. Plat-E cells packaged the libraries, and Ba/F3 cells were infected with the produced retrovirus. Following drug selection, the cells were exposed for 2 weeks to repotrectinib or ensartinib, and resistant mutations were identified by sequencing the alive clones. (b) The targeted amino acid sequence for error-prone PCR (GenBank: AB663645, 1850-2383). Major ALK-TKI resistance mutations were identified, and all of them were included in the targeted region for error-prone PCR.

At first, error-prone PCR was performed against the pMXs vector, which contained cDNA of EML4-ALK variant 1 with G1202R or I1171N mutation.

All of resistant mutations observed in the clinic were covered by inducing mutations at random with error-prone PCR into the sequence encoding the ALK kinase domain (534 bp) (Fig.2b). In this method, the number of PCR cycles and the amount of DNA templates can be adjusted to control the frequency of mutations. To detect the resistant mutations caused by a single base substitution, we aimed to set the mutation frequency to 0.2 mutations/copy. According to Poisson’s distribution calculation, the percentage of PCR products with two or more mutations against all PCR products would be approximately 10 times lower than that of products with a single mutation in this condition. Indeed, NGS analysis revealed that the constructed libraries exhibited a mutation frequency of 0.24 mutations/copy.

Then, the PCR products were combined with pMXs vector arms and then introduced into E. coli. cDNA mutant library was constructed from 7 × 10^5^ colonies. NGS analysis of the constructed libraries (300 bp) revealed a variety of patterns of base substitutions in the targeted region (Fig.S2a). According to the NGS analysis, the lowest mutation pattern frequency (A > C, T > G) occurred in 3.6% of the PCR products. If this base substitution occurs equally in all adenine and thymine bases (130 bp), 2.77 × 10^−2^ (%) of PCR products will harbor a mutation in the specific adenine or thymine base. According to Poisson’s distribution, 1.2 × 10^4^ clones are required to obtain the specific mutant. Thus, although these mutation patterns were somewhat biased (Fig.S2b), possible mutations appeared to be included due to the library size (7 × 10^5^). Biased mutation patterns (Fig.S2c) may reflect the targeted region’s GC content (58%).

At last, the cDNA libraries were transfected into Plat-E cells and packaged into a retrovirus, and then, 3 × 10^7^ cells were incubated with the virus. The observation of cell growth after the addition of puromycin revealed that their infection efficiency was greater than 10% (Fig.S2d). Given that the cDNA mutant library was approximately 7 × 10^5^ colonies in size and that more than 3 × 10^6^ cells were infected, every possible mutation was likely to be included in the established Ba/F3 cell libraries.

### Prediction of repotrectinib- or ensartinib-resistant mutations

To confirm the validity of the method for the error-prone PCR based method for predicting resistant mutations, we predicted alectinib-resistant mutations. After exposure to 50 nM alectinib for a week, the proliferated Ba/F3 cells expressing the mutant library were analyzed to identify resistant mutations by NGS. As a result, we identified the alectinib-resistant mutations I1171N/T/S, V1180L, L1196M, L1198F, and G1202R. (Fig.3a). Most of alectinib-resistant mutations observed in the clinic were detected by using our system^9,12^.

Next, we practically predicted resistant mutations against repotrectinib or ensartinib. The established mutant library expressing Ba/F3 cells were exposed to repotrectinib or ensartinib for 2 weeks. The concentrations of repotrectinib and ensartinib were set at 1.5 μM, 800 nM, respectively, to allow only resistant mutants to proliferate. To obtain a single clone of the resistant mutant, we minimized the number of cells seeded into each well of a 96-well plate so that the proliferation was observed in no more than 15 wells. In this situation, these cells probably proliferated from a single clone. We predicted several resistant compound mutations against these TKIs by sangar sequencing after 2 weeks of exposure to each inhibitor (Fig.3b,c). As a result, we predicted L1196M +, F1174C/L/I +, E1129V/K +, and C1156Y + G1202R as repotrectinib-resistant compound mutations (Fig.3b). In the clinic, the compound mutations L1196M + G1202R and F1174L + G1202R were referred to as lorlatinib-resistant mutations^13,24^. C1156Y and E1129V single mutations have been observed in patients who relapsed after crizotinib treatment^25,26^. Also, these resistant mutations were discovered by the bulk culture’s NGS analysis (Fig.S3a). On the other hand, we predicted C1156Y +, L1256F +, and E1129V/K + I1171N mutations as ensartinib-resistant compound mutations (Fig3c). Of them, C1156Y + I1171N compound mutation has been observed after alectinib and ceritinib treatments^9^. In addition to these resistant mutations, the bulk culture’s NGS analysis revealed H1124R + and F1174I + I1171N as minor resistant mutations (Fig.S3b).

**Fig 3.**
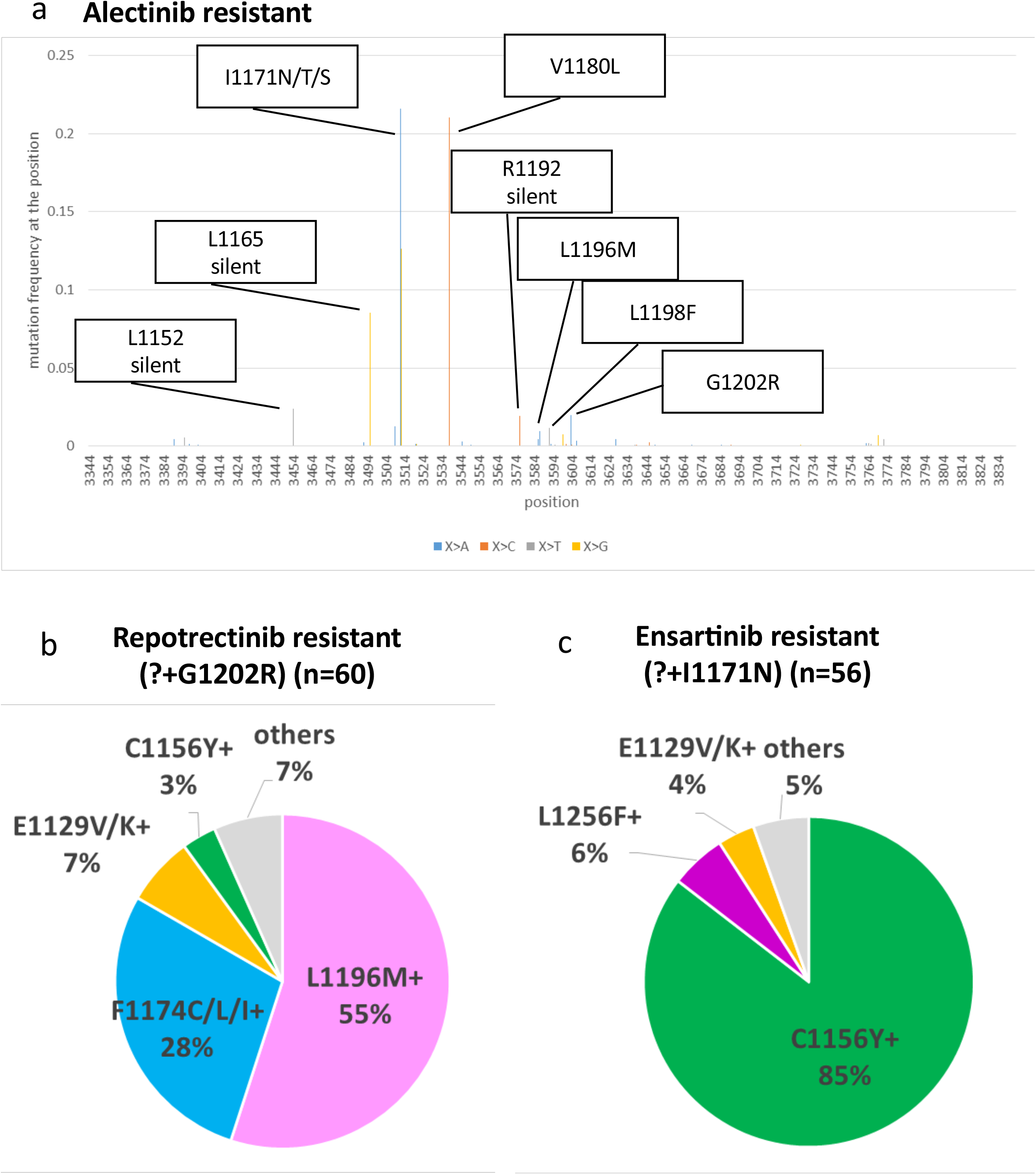
Predicted resistant mutations against ALK-TKIs. (a) Predicted alectinib-resistant mutations. 3 × 10^6^ mutant library expressing Ba/F3 cells were exposed to alectinib (50 nM) for 1 week, and genomic DNA was extracted. PCR amplicons were produced from the DNA and sequenced by NGS. (b) Predicted repotrectinib-resistant mutations (n = 60). Mutant libraries expressing Ba/F3 cells were exposed to repotrectinib (1.5 μM) for 2 weeks. Others contain G1202R + I1171T and L1196M + T1151M + G1202R. (c) Predicted ensartinib-resistant mutations (n = 55). For 2 weeks, mutant library expressing Ba/F3 cells were exposed to ensartinib (800 nM). I1171N + H1124R, I1171N + A1200R, and I1171N + L1196M are among the others.

### Sensitivity of the predicted resistant mutants to other TKIs

To determine potential therapeutic strategies following repotrectinib or ensartinib treatment, we assessed the sensitivity of acquired repotrectinib- and ensartinib-resistant mutants to various TKIs. First, we established Ba/F3 cells expressing EML4-ALK with predicted repotrectinib- or ensartinib-resistant compound mutation, and then, we assessed the sensitivity using these mutants. Almost all of these mutants were resistant to alectinib, except the L1256F+ I1171N mutant (Fig.4c,d), which was consistent with previous research^15^. Alectinib inhibited phosphorylation of ALK with the L1256F + I1171N mutation according to immunoblotting (Fig.4e).

**Fig 4.**
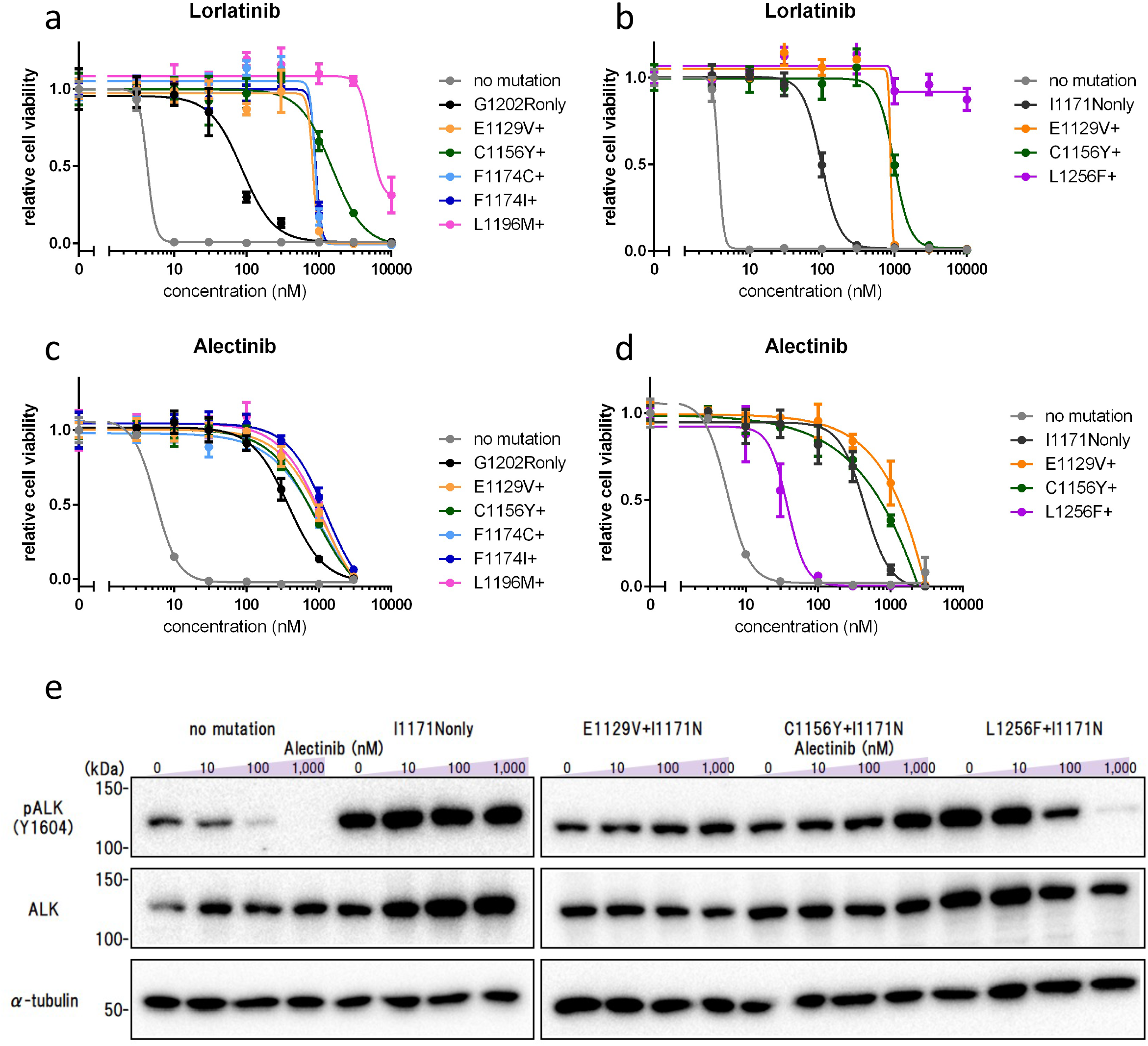
Sensitivity of predicted mutants against lorlatinib and alectinib. (a–d) Sensitivity evaluation of repotrectinib- or ensartinib-resistant mutants against lorlatinib (a,b) and alectinib (c,d). Each inhibitor was given to Ba/F3 cells expressing EML4-ALK variant 1 with a repotrectinib- or ensartinib-resistant compound mutation for 72 h. CCK-8 and absorbance at 450 nm were used to assess cell viability. (e) Immunoblotting evaluation of phosphorylated ALK suppression in ensartinib-resistant mutants by alectinib. Alectinib was given to Ba/F3 cells expressing EML4-ALK v1 with each resistant mutation for 3 h. Next, immunoblotting was used to detect the indicated proteins in cell lysates.

TPX-0131 is a next-generation ALK-TKI that is potent to lorlatinib-resistant compound mutations such as L1196M + G1202R^27^. In addition to the L1196M + G1202R mutation, we discovered that repotrectinib-resistant mutants with the C1156Y + and E1129V + G1202R mutations were sensitive to TPX-0131 (Fig.5a). Conversely, the F1174C/I + G1202R repotrectinib-resistant mutant and all of ensartinib-resistant mutants were resistant to TPX-0131 (Fig.5a,b). Immunoblotting revealed that phosphorylation of ALK with L1196M +, C1156Y +, or E1129V + G1202R mutation was sufficiently suppressed by TPX-0131 (Fig.5c).

**Fig 5.**
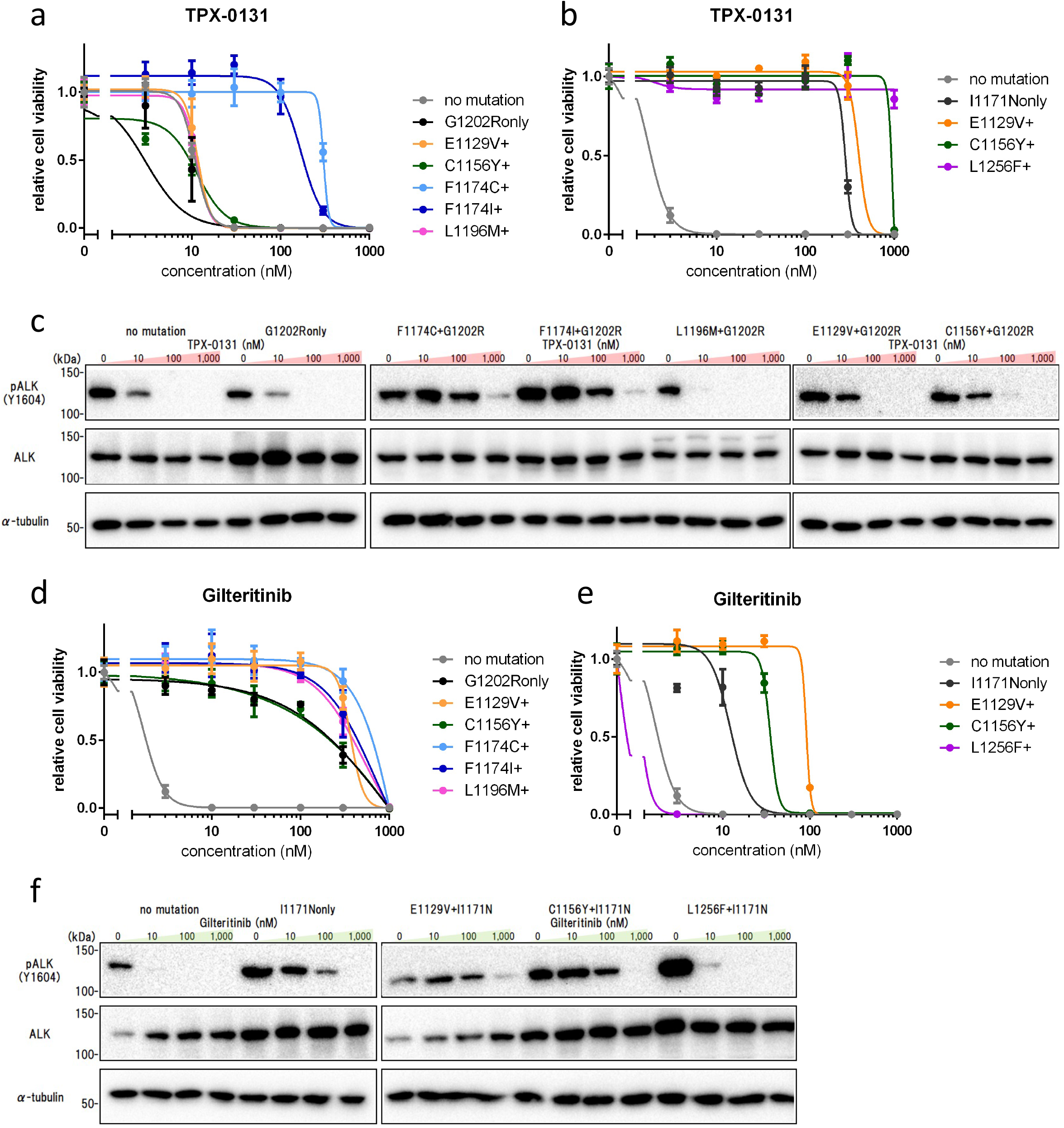
Sensitivity of predicted mutants against TPX-0131 and gilteritinib. (a–c) Sensitivity evaluation of repotrectinib- or ensartinib-resistant mutants against TPX-0131 by CCK-8 (a,b) and immunoblotting (c). Ba/F3 cells expressing EML4-ALK variant 1 with a repotrectinib- or ensartinib-resistant compound mutation were exposed to each inhibitor for 72 h for the CCK-8 viability assay. Ba/F3 cells expressing EML4-ALK v1 with each resistant mutation were exposed to each inhibitor for 3 hours before immunoblotting. (d–f) CCK-8 (d,e) and immunoblotting were used to assess the sensitivity of repotrectinib- or ensartinib-resistant mutants to gilteritinib (f).

Since the previous study shows that adaphostin, a BCR-ABL inhibitor, was potent to L1196M + G1202R mutants^15^ and gilteritinib, a FLT3 inhibitor, was potent to various I1171N compound mutants including L1256F + I1171N^16^, we assessed the potency of each inhibitor against the acquired mutants. The cell viability assay revealed that no repotrectinib-resistant mutants were sensitive to gilteritinib (Fig.5d), and none of the acquired mutants were sensitive to adaphostin (Fig.S4c,d). Conversely, ensartinib-resistant mutants C1156Y + I1171N and L1256F + I1171N were sensitive to gilteritinib (Fig.5e), and immunoblotting revealed that phosphorylation of ALK with these mutations was well suppressed (Fig.5f).

## Discussion

Lung cancer is the most commonly diagnosed cancer and the leading cause of cancer-related death around the world^28^. NSCLC accounts for approximately 85% of all lung cancer diagnoses^29^. Several oncogenic driver genes have been identified in NSCLC, and molecular-targeted drugs have been developed to improve the disease’s prognosis^30,31^. ALK rearrangement has been observed in approximately 4% of patients with NSCLC^1,2^, and different ALK-TKIs have been used in the clinic. Alectinib, a second-generation ALK-TKI, in particular, demonstrated extremely long progression-free survival than other standard chemotherapy^8^, and is used as the first-line therapy. The third-generation ALK-TKI, lorlatinib, is potent to alectinib-resistant mutants^10,11^, and usage of lorlatinib as the first-line treatment has been approved recently due to its efficacy^17^. However, patients will inevitably relapse due to the appearance of resistant mutants even after lorlatinib treatment^12,13^. Thus developing valuable sequential therapeutic strategies that maximize the efficacy of ALK-TKIs-treatments is critical.

To establish other effective second-line therapeutic strategies than lorlatinib treatment after the first-line alectinib treatment, prediction of compound resistant mutations is required because many patients relapse due to secondary mutations in ALK kinase domain after ALK-TKIs treatment^9,12,13^. In this report, we established a controllable and comprehensive method for predicting resistant mutations using error-prone PCR (Fig.2). Because the contents and size of mutant libraries can be controlled in this method, the prediction of resistant mutations appears to be more comprehensive than the conventional method based on ENU mutagenesis^13,15^. Moreover, the off-target mutations are never occurred in the error-prone PCR method, which differs from ENU mutagenesis. Indeed, the control experiment using alectinib predicted almost all of alectinib-resistant mutations discovered in the clinic.

Previous research revealed that repotrectinib was potent to alectinib-resistant G1202R mutant^22^; however, whether repotrectinib and ensartinib are potent to alectinib-resistant I1171N mutant has not been clarified. We assessed I1171N mutant to be sensitive to these inhibitors *in vitro* (Fig.1c–d). As a result, ensartinib was found to be potent against the I1171N mutant, and these two inhibitors has possibility to be second-line options after alectinib treatment. Even though those inhibitors can be applicable after the failure of alectinib treatment, the additional resistant mutations against them will be generated. Therefore, we predicted resistant compound mutations against virtual second-line treatment with repotrectinib or ensartinib after the first-line alectinib treatment as a model study (Fig.3).

Then we assessed the sensitivity of the predicted resistant mutants against several inhibitors (Fig.4,5) and showed that most of the mutants were resistant to approved ALK-TKIs as were the lorlatinib-resistant mutations. Therefore other TKIs should be used to treat patients with ALK-positive NSCLC who have become resistant to repotrectinib or ensartinib (Fig.6). L1196M + G1202R, a major repotrectinib-resistant mutation, is also discovered to be a lorlatinib-resistant mutation^13^. This mutant is highly resistant to all approved ALK-TKIs; however, we confirmed that a novel ALK-TKI, TPX-0131, was potent to this mutant^27^. If the efficacy and safety of TPX-0131 were demonstrated in a current clinical trial, this inhibitor would be an excellent treatment option for repotrectinib-resistant patients with the L1196M + G1202R mutation. However, we showed that TPX-0131 was not potent to F1174C/I + G1202R and ensartinib-resistant mutants. Conversely, brigatinib may be effective for F1174C/I + G1202R mutants because its IC50 for F1174C/I + G1202R is slightly lower than that of the other compound mutation containing G1202R^15,32^.

**Fig 6.**
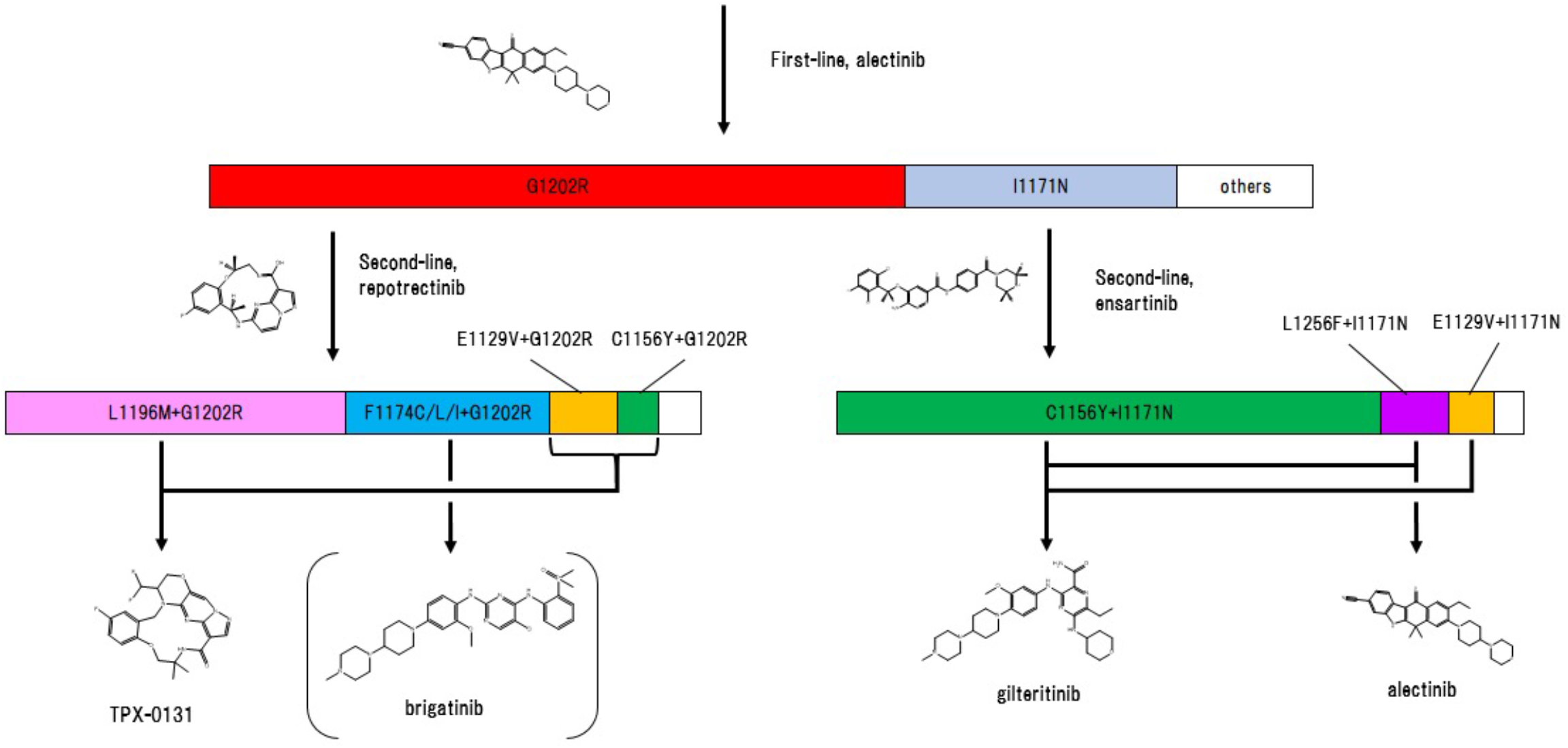
Possible therapeutic strategies after repotrectinib or ensartinib treatment. Repotrectinib is available for G1202R harboring ALK + NSCLC after alectinib treatment. Some patients may relapse as a result of another L1196M mutation. In such a case, TPX-0131 will be a viable treatment option. Additional F1174C/L/I mutations will occasionally be observed, which will possibly be treated with brigatinib. Ensartinib is a treatment option for I1171N patients with ALK-positive NSCLC. The majority of patients may relapse due to an additional C1156Y mutation. In this case, gilteritinib will be potent. Rarely recurrence will be caused by additional L1256F mutations, which can be treated with alectinib or gilteritinib. Each length of compound mutation reflects the ratio of the respective number of obtained mutant clones depicted in Fig.3b–c.

According to our predictive screening using ensartinib, C1156Y + I1171N would be identified as the most frequent ensartinib-resistant mutation. Although this mutant was resistant to all of ALK-TKIs to be evaluated, gilteritinib was potent to it. If gilteritinib was approved for the treatment of patients with ALK-positive NSCLC, this therapeutic strategy would be effective. We identified L1256F + I1171N as a minor ensartinib-resistant mutation. This mutant would be overcome by alectinib or gilteritinib treatment^15,16^. Also, these resistant mutations predicted in our study were identified as lorlatinib-resistant mutations in previous studies (Table.1)^13,15^. Our findings highlighted the importance of effective therapeutic strategies for these hotspot mutations. In addition to current sequential therapies, combinational therapy using ALK-TKIs and other inhibitors at the same time must be investigated to completely prevent the generation of hotspot resistant mutants.

In conclusion, we developed an error-prone PCR method for predicting drug-resistant mutations. As a model study, we predicted repotrectinib or ensartinib-resistant mutations and proposed that repotrectinib and ensartinib could be available for the alternative second-line usage against patients who do not apply to treatment with lorlatinib. Lorlatinib is currently approved for the first-line treatment. A single course of lorlatinib treatment may provide more clinical benefit than a series of treatments that would accumulate genetic changes such as ALK compound mutations. Nevertheless, a variety of clinical strategies must be established for each patient to receive the best linical output while maintaining a high-quality of life.

**Table 1.**
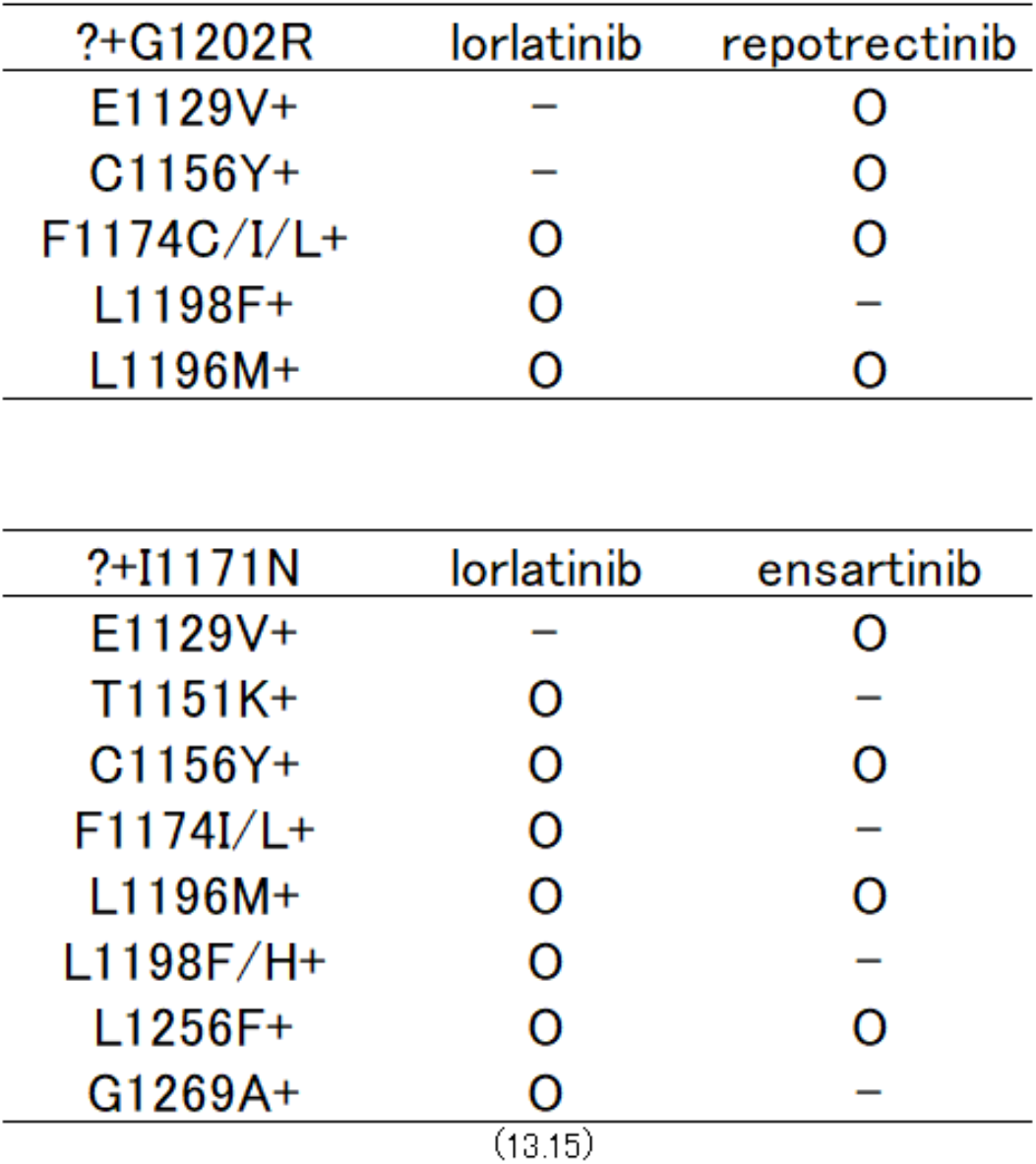
Predicted resistant mutations in this method and in previous studies.

| ?+G1202R | lorlatinib | repotrectinib |
| --- | --- | --- |
| E1129V+ | – | O |
| C1156Y+ | – | O |
| F1174C/I/L+ | O | O |
| L1198F+ | O | – |
| L1196M+ | O | O |

| ?+I1171N | lorlatinib | ensartinib |
| --- | --- | --- |
| E1129V+ | – | O |
| T1151K+ | O | – |
| C1156Y+ | O | O |
| F1174I/L+ | O | – |
| L1196M+ | O | O |
| L1198F/H+ | O | – |
| L1256F+ | O | O |
| G1269A+ | O | – |

Resistant mutations against lorlatinib predicted with the ENU method^13,15^, or against repotrectinib and ensartinib predicted with error-prone PCR method, are indicated as O. Mutations were predicted by exposure to 300, 600, 1,000 nM lorlatinib, 1.5 μM repotrectinib or 800 nM ensartinib, respectively.

## Acknowledgement

The authors would like to thank Enago (www.enago.jp) for the English launguage review.

We thank Prof. T. Kitamura (Institute of Medical Science, The University of Tokyo, Tokyo, Japan) for kindly providing the Plat-E cell line.

We appreciate to Dr. Fujimoto (Japan Biologycal informatics Consortium (JBiC), Tokyo, Japan) for the experimental direction.

## Funding

This study was partly supported by the Fukushima Translational Research Project program.

## Authors’ contributions

K.S. conceived and designed this study. Y.D, H.T., A.N. performed the experiments and analyzed the data. Y.D and K.S. interpreted the data and wrote the manuscript. All the authors reviewed and edited the manuscript.

## Conflict of interest

The authors declare no conflict of interest.

## Notes

### Competing Interest Statement

The authors have declared no competing interest.

## References

1. Jordan, E. J. et al. Prospective comprehensive molecular characterization of lung adenocarcinomas for efficient patient matching to approved and emerging therapies. Cancer Discov. 7, 596–609 (2017).

2. Wang, S. et al. Assessment of nine driver gene mutations in surgically resected samples from patients with non-small-cell lung cancer. Cancer Manag. Res. 12, 4029–4038 (2020).

3. Soda, M. et al. Identification of the transforming EML4-ALK fusion gene in non-small-cell lung cancer. Nature 448, 561–566 (2007).

4. Morris, S. W. et al. Fusion of a kinase gene, ALK, to a nucleolar protein gene, NPM, in non-Hodgkin’s lymphoma. Science 263, 1281–1284 (1994).

5. Bischof, D., Pulford, K., Mason, D. Y. & Morris, S. W. Role of the nucleophosmin (NPM) portion of the non-Hodgkin’s lymphoma-associated NPM-anaplastic lymphoma kinase fusion protein in oncogenesis. Mol. Cell. Biol. 17, 2312–2325 (1997).

6. Duyster, J., Bai, R. Y. & Morris, S. W. Translocations involving anaplastic lymphoma kinase (ALK). Oncogene 20, 5623–5637 (2001).

7. Solomon, B. J. et al. First-line crizotinib versus chemotherapy in ALK-positive lung cancer. N. Engl. J. Med. 371, 2167–2177 (2014).

8. Peters, S. et al. Alectinib versus crizotinib in untreated ALK-positive non–small-cell lung cancer. N. Engl. J. Med. 377, 829–838 (2017).

9. Gainor, J. F. et al. Molecular mechanisms of resistance to first-and second-generation ALK inhibitors in ALK-rearranged lung cancer. Cancer Discov. 6, 1118–1133 (2016).

10. Shaw, A. T. et al. Lorlatinib in non-small-cell lung cancer with ALK or ROS1 rearrangement: An international, multicentre, open-label, single-arm first-in-man phase 1 trial. Lancet Oncol. 18, 1590–1599 (2017).

11. Zou, H. Y. et al. PF-06463922, an ALK/ROS1 inhibitor, overcomes resistance to first and second generation ALK inhibitors in preclinical models. Cancer Cell. 28, 70–81 (2015).

12. Dagogo-Jack, I. et al. Treatment with next-generation ALK inhibitors fuels plasma ALK mutation diversity. Clin. Cancer Res. 25, 6662–6670 (2019).

13. Yoda, S. et al. Sequential ALK inhibitors can select for lorlatinib-resistant compound ALK mutations in ALK-positive lung cancer. Cancer Discov. 8, 714–729 (2018).

14. Shaw, A. T. et al. Resensitization to crizotinib by the Lorlatinib ALK resistance mutation L1198F. N. Engl. J. Med. 374, 54–61 (2016).

15. Okada, K. et al. Prediction of ALK mutations mediating ALK-TKIs resistance and drug re-purposing to overcome the resistance. EBiomedicine. 41, 105–119 (2019).

16. Mizuta, H. et al. Gilteritinib overcomes lorlatinib resistance in ALK-rearranged cancer. Nat. Commun. 12, 1261 (2021).

17. Shaw, A. T. et al. First-line Lorlatinib or crizotinib in advanced ALK -Positive lung cancer. N. Engl. J. Med. 383, 2018–2029 (2020).

18. Bauer, D. C., McMorran, B. J., Foote, S. J. & Burgio, G. Genome-wide analysis of chemically induced mutations in mouse in phenotype-driven screens. BMC Genomics. 16, 866 (2015).

19. Gibson, D. G. et al. Enzymatic assembly of DNA molecules up to several hundred kilobases. Nat. Methods. 6, 343–345 (2009).

20. Bolger, A. M., Lohse, M. & Usadel, B. Trimmomatic: A flexible trimmer for Illumina sequence data. Bioinformatics. 30, 2114–2120 (2014).

21. Kim, D., Paggi, J. M., Park, C., Bennett, C. & Salzberg, S. L. Graph-based genome alignment and genotyping with HISAT2 and HISAT-genotype. Nat. Biotechnol. 37, 907–915 (2019).

22. Drilon, A. et al. Repotrectinib (Tpx-0005) is a next-generation ros1/trk/alk inhibitor that potently inhibits ros1/trk/alk solvent-front mutations. Cancer Discov. 8, 1227–1236 (2018).

23. Horn, L. et al. Ensartinib (X-396) in ALK-positive non–small cell lung cancer: Results from a first-in-human phase I/II, multicenter study. Clin. Cancer Res. 24, 2771–2779 (2018).

24. Recondo, G. et al. Diverse resistance mechanisms to the third-generation ALK inhibitor lorlatinib in ALK-rearranged lung cancer. Clin. Cancer Res. 26, 242–255 (2020).

25. Tan, D. S. et al. Genetic landscape of patients with ALK-rearranged non–small-cell lung cancer (NSCLC) and response to ceritinib in ASCEND-1 study. Lung Cancer. 163, 7–13 (2022).

26. Choi, Y. L. et al. EML4-ALK mutations in lung cancer that confer resistance to ALK inhibitors. N. Engl. J. Med. 363, 1734–1739 (2010).

27. Murray, B. W. et al. TPX-0131, a potent CNS-penetrant, next-generation inhibitor of wild-type ALK and ALK-resistant mutations. Mol. Cancer Ther. 20, 1499–1507 (2021).

28. Bray, F. et al. Global cancer statistics 2018: GLOBOCAN estimates of incidence and mortality worldwide for 36 cancers in 185 countries. CA Cancer J. Clin. 68, 394–424 (2018).

29. Oser, M. G. et al. Transformation from non-small-cell lung cancer to small-cell lung cancer: Molecular drivers and cells of origin. Lancet Oncol. 16, e165–e172 (2015).

30. Halliday, P. et al. Emerging targeted therapies for the treatment of non-small cell lung cancer. Curr. Oncol. Rep. 21, 21 (2019).

31. Yuan, M. et al. The emerging treatment landscape of targeted therapy in non− small-cell lung cancer. Signal Transduct. Target. Ther. 4, 61 (2019).

32. Song, X. et al. Two novel strategies to overcome the resistance to ALK tyrosine kinase inhibitor drugs: Macrocyclic inhibitors and proteolysis-targeting chimeras. Med. 2, 341–350 (2021).

