## Supplemental Figures for "Prediction of Resistant Mutations against Upcoming ALK-TKIs, Repotrectinib (TPX-0005) and Ensartinib (X-396)"

**Legends to supplementary figures**

**Fig.S1 Suppression of phosphorylation of ALK by exposure to repotrectinib or ensartinib**

(a,b) Immunoblotting evaluation of phosphorylated ALK reduction in alectinib-resistant mutants by repotrectinib or ensartinib. Ba/F3 cells expressing EML4-ALK v1 with G1202R or I1171N were exposed to repotrectinib or ensartinib for 3 h. Immunoblotting was used to detect the indicated proteins in cell lysates.

**Fig.S2 Evaluation of cDNA mutant libraries**

(a) Mutation heatmap analyzed by NGS. Left axis reveals variations of mutations and their mutation frequency at the position. Right axis shows the total amount of high-quality reads (Qscore > 30). (b,c) Patterns of induced mutations by error-prone PCR (n = 3). (d) Cell growth observation after puromycin addition for calculating infection efficacy. Cell concentration in 0 h was $5 \times 10^5$ (cells/mL). Alive cells were counted at 41, 48, 65, 72 h after puromycin addition and drew regression lines. Confidence of determination was 0.973 (I1171N) or 0.965 (G1202R) and predicted initial infected cells concentration was $0.315 \times 10^5$ (I1171N) or 0.498 × 10e5 (G1202R). As a result, infection efficiency was calculated as 15.8% (I1171N) or 24.9% (G1202R).

**Fig.S3 Predicted resistant mutations against repotrectinib or ensartinib by NGS sequencing**

(a,b) Predicted repotrectinib- or ensartinib-resistant mutations. $3 \times 10^6$ of mutant library expressing Ba/F3 cells were exposed to repotrectinib (1.5 μM) or ensartinib (800 nM) for 1 week, and genomic DNA was extracted. PCR amplicons were produced from the DNA and sequenced by NGS.

**Fig.S4 Sensitivity of predicted mutants against crizotinib and adaphostin.**

(a–d) Sensitivity evaluation of repotrectinib- or ensartinib-resistant mutants against crizotinib (a,b) and adaphostin (c,d). Ba/F3 cells expressing EML4-ALK variant 1 with a repotrectinib- or ensartinib-resistant compound mutation were exposed to each inhibitor for 72 h. Cell viability swas assessed by CCK-8 and absorbance at a 450 nm wave length.

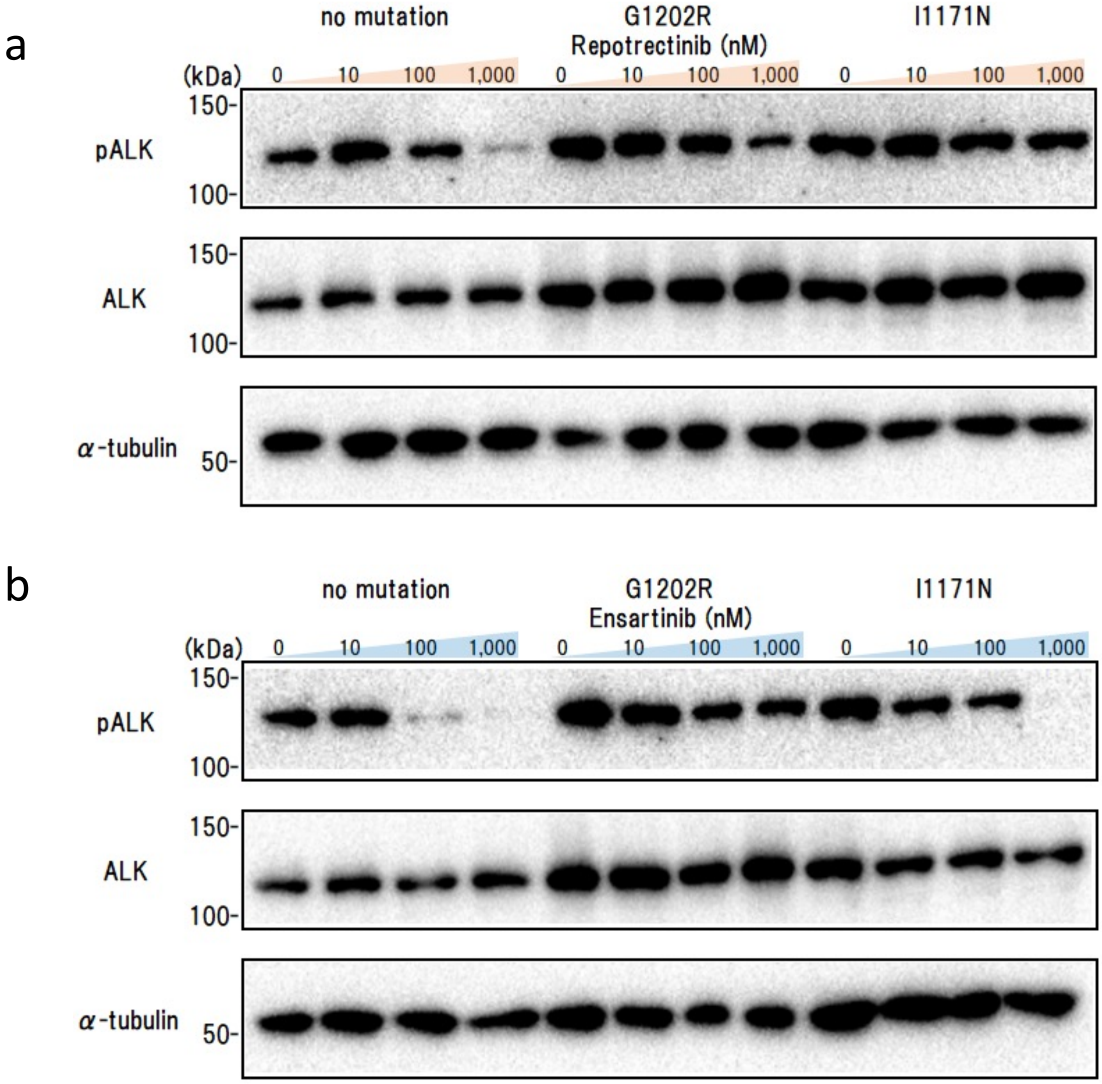
a
no mutation
G1202R
I1171N
Repotrectinib (nM)
(kDa)
0
10
100
1,000
150-
100-
pALK
ALK
α-tubulin
50-
b
no mutation
G1202R
I1171N
Ensartinib (nM)

Fig.S1

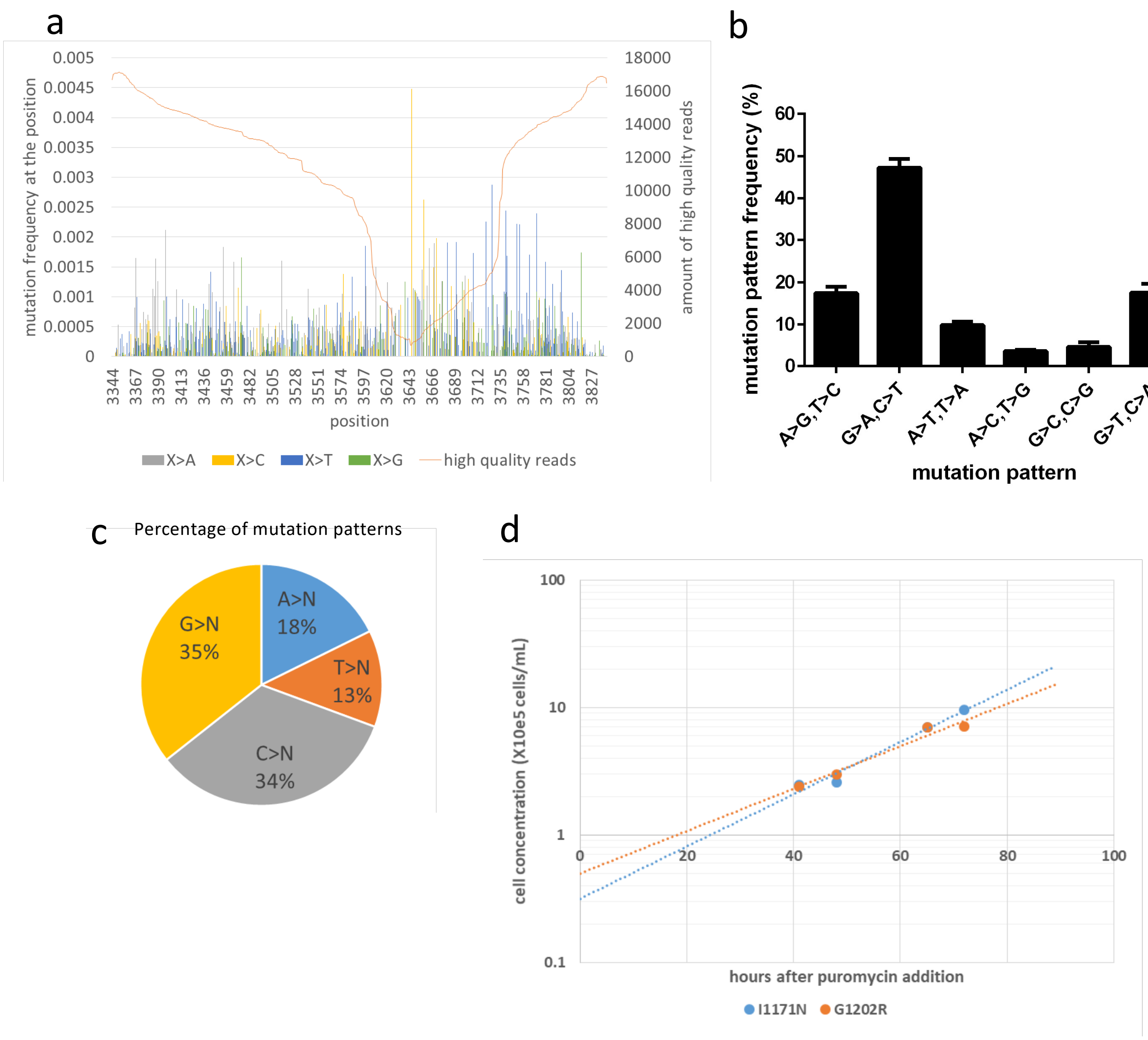
a
mutation frequency at the position
amount of high quality reads
position
X>A
X>C
X>T
X>G
high quality reads
b
mutation pattern frequency (%)
A>G,T>C
G>A,C>T
A>T,T>A
A>C,T>G
G>C,C>G
G>T,C>A
mutation pattern
c
Percentage of mutation patterns
A>N 18%
T>N 13%
C>N 34%
G>N 35%
d
cell concentration (X10e5 cells/mL)
hours after puromycin addition
I1171N
G1202R

Fig.S2

a **Repotrectinib resistant(?+G1202R)**

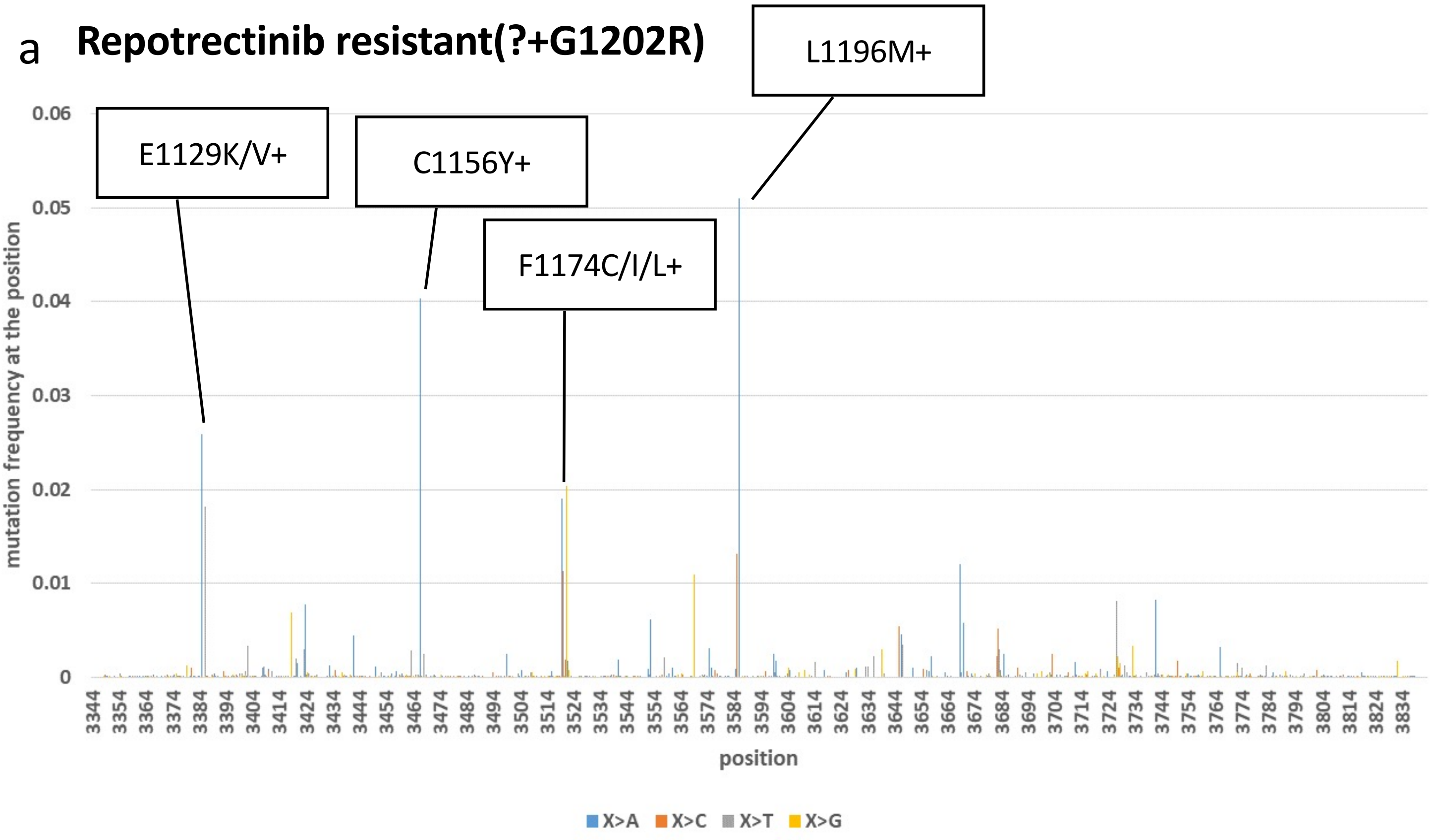


b **Ensartinib resistant (?+I1171N)**

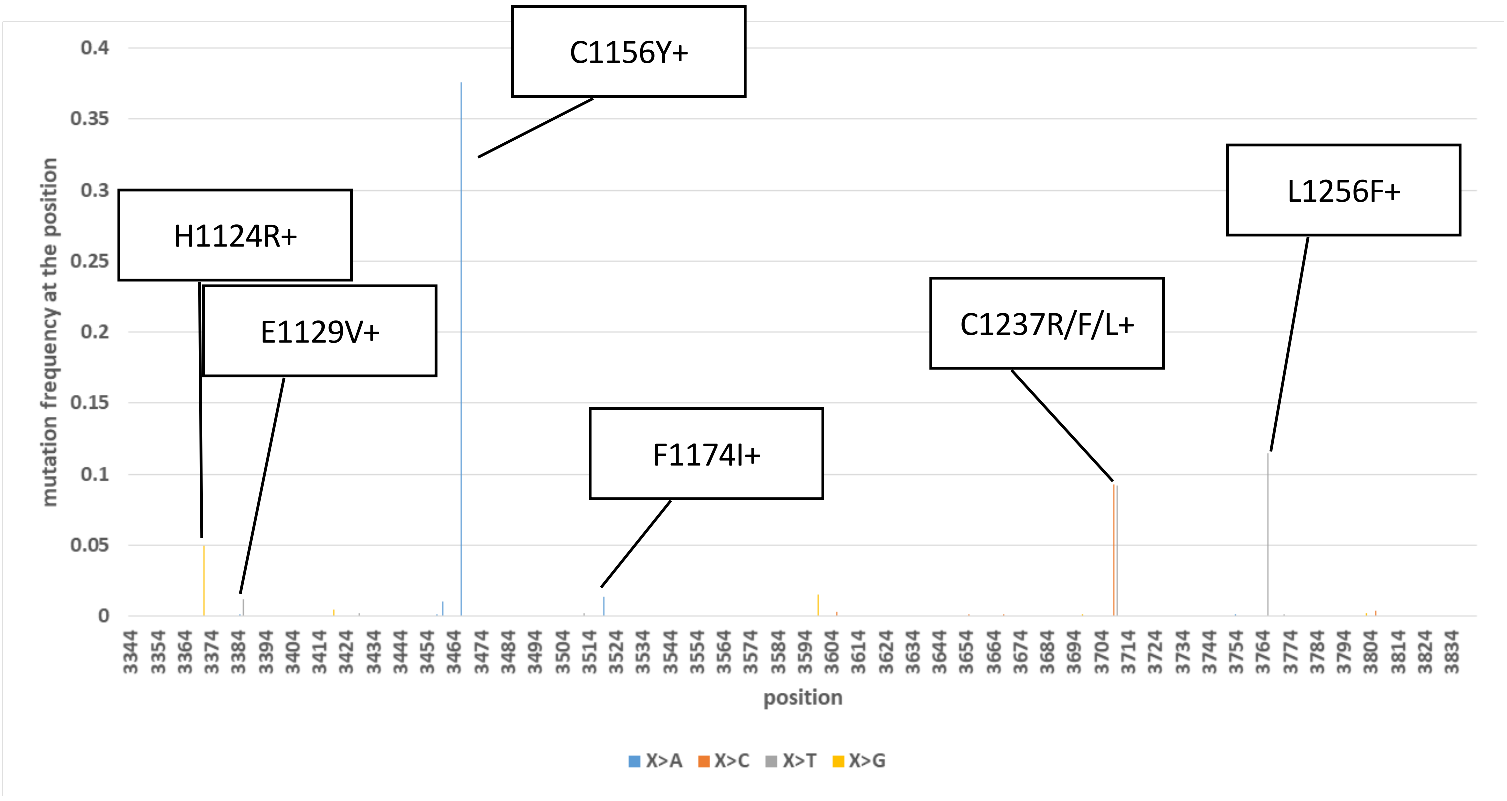


Fig.S3

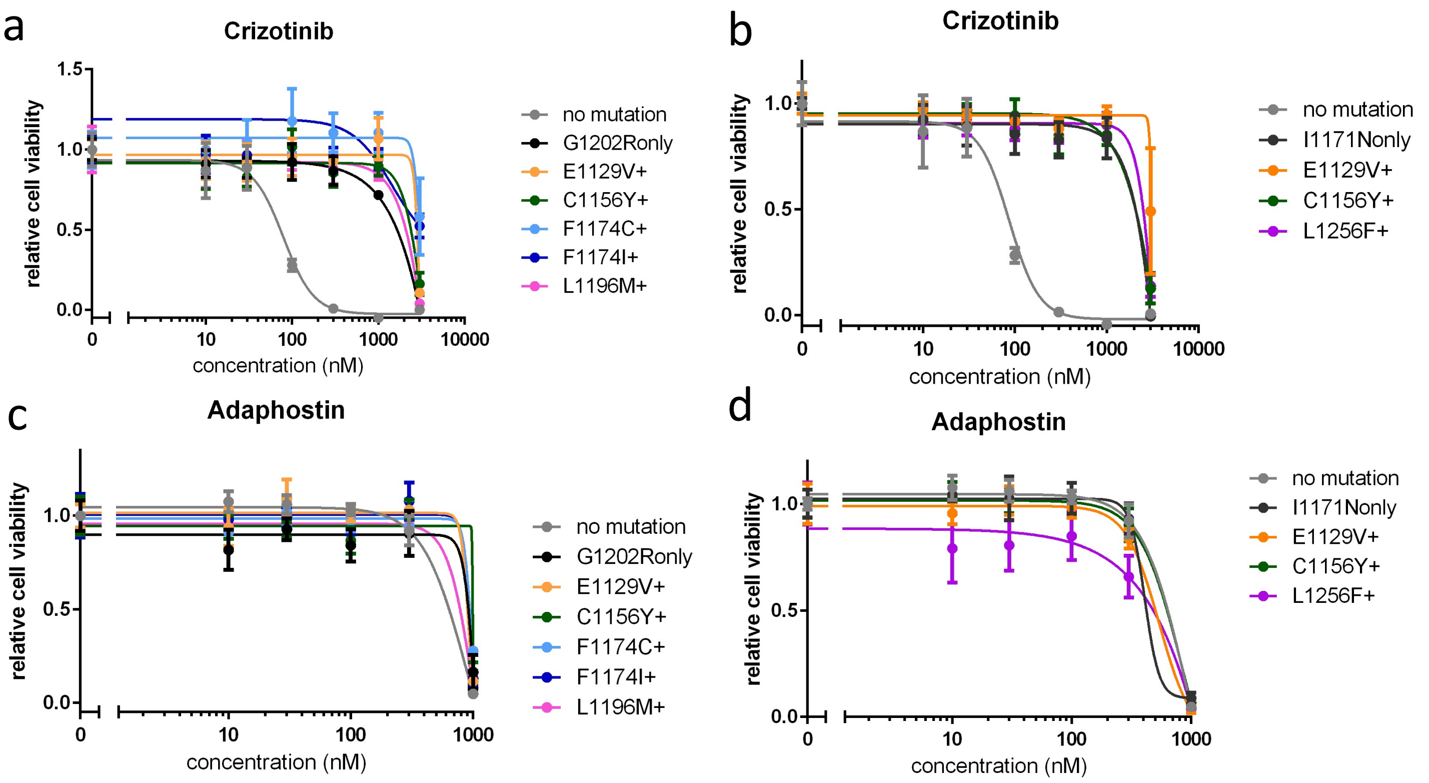

a
Crizotinib
relative cell viability
1.5
1.0
0.5
0.0
0
10
100
1000
10000
concentration (nM)
no mutation
G1202Ronly
E1129V+
C1156Y+
F1174C+
F1174I+
L1196M+
b
Crizotinib
relative cell viability
1.0
0.5
0.0
0
10
100
1000
10000
concentration (nM)
no mutation
I1171Nonly
E1129V+
C1156Y+
L1256F+
c
Adaphostin
relative cell viability
1.0
0.5
0.0
0
10
100
1000
concentration (nM)
no mutation
G1202Ronly
E1129V+
C1156Y+
F1174C+
F1174I+
L1196M+
d
Adaphostin
relative cell viability
1.0
0.5
0.0
0
10
100
1000
concentration (nM)
no mutation
I1171Nonly
E1129V+
C1156Y+
L1256F+


Fig.S4
